# LRH-1 Mitigates Intestinal Inflammatory Disease by Maintaining Epithelial Homeostasis and Cell Survival

**DOI:** 10.1101/314302

**Authors:** James R. Bayrer, Hongtao Wang, Roy Nattiv, Miyuki Suzawa, Hazel S. Escusa, Robert J. Fletterick, Ophir D. Klein, David D. Moore, Holly A. Ingraham

**Author notes:** These authors contributed equally (James R. Bayrer, Hongtao Wang).

## Abstract

Epithelial dysfunction and loss of intestinal crypts are defining features of inflammatory bowel disease (IBD). However, current therapies primarily target the immune system and not the epithelium. The nuclear receptor LRH-1 encoded by *Nr5a2* is expressed in intestinal epithelium and is thought to contribute to epithelial renewal. Here we investigate how loss and gain of LRH-1 impacts the intestinal epithelium in healthy and inflammatory conditions. Knocking out LRH-1 in murine intestinal organoids reduces Notch signaling, increases crypt cell death and weakens the epithelial barrier. Loss of LRH-1 also distorts the cellular composition of the epithelium, resulting in an expansion of Paneth and goblet cells, and a decrease in enteroendocrine cells. Human LRH-1 (hLRH-1) not only rescues epithelial integrity, but when overexpressed, mitigates inflammatory damage in mouse and human intestinal organoids, including those from IBD patients. Finally, hLRH-1 greatly reduces disease severity in a mouse model of T cell-mediated colitis. Together with the failure of a ligand-incompetent hLRH-1 mutant to protect against TNFα-damage, these findings provide compelling evidence that hLRH-1 mediates epithelial homeostasis and is an attractive target for intestinal disease.

---

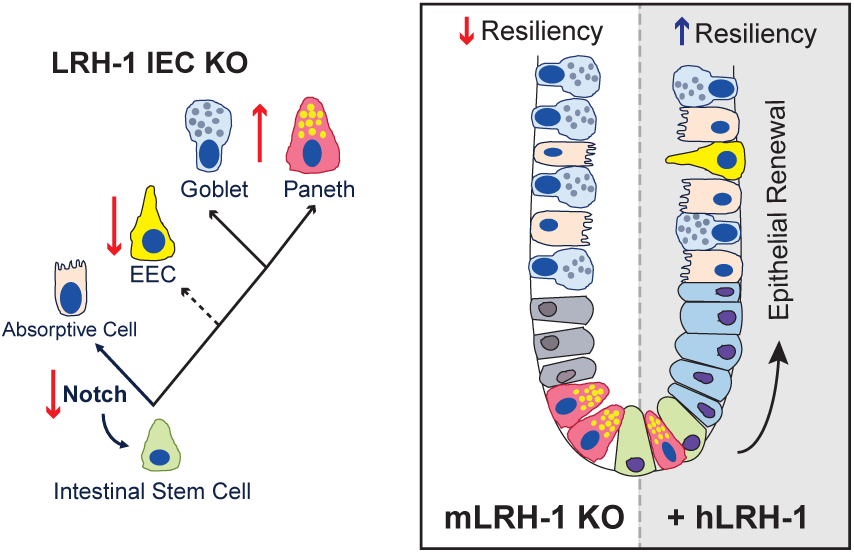

Inflammatory bowel disease (IBD) is a chronic disorder that is characterized by bouts of intense gastrointestinal inflammation, ultimately resulting in destruction of the epithelial lining of the gut^1^. Although defects in genes expressed in the gut epithelium have been associated with IBD^2,3^, the contribution of the epithelium to this disease remains understudied, particularly in comparison to the intensive interrogation of the immune component. However, the recent establishment of mouse and human intestinal organoids has provided an excellent experimental platform to explore intrinsic epithelial defects in patients and mouse models with disease^4,5^.

An important regulatory factor for intestinal epithelia is Liver Receptor Homolog 1 (LRH-1, NR5A2). This nuclear receptor has been shown to be expressed in intestinal crypts, where intestinal stem cells (ISCs) reside^6^, and where it contributes to epithelial renewal by potentiating WNT/b-catenin signaling^6,7,8^. Recent GWAS meta-analyses of IBD patients found a significant association between LRH-1 and a form of IBD called ulcerative colitis^9,10^. Animal studies using heterozygous (*Lrh-1^+/−^*) or conditional knock out (*Lrh-1^fl/fl^; VilCre-Ert2*) did not note any apparent epithelial defects at baseline, but did report a defect in epithelial proliferation and susceptibility to colitis^6,11^. Interestingly, the elimination of LRH-1 in mouse intestine and human colon cancer cell lines resulted in decreased glucocorticoid production^12,13,14^, which has the potential to lead to the kind of increased intestinal inflammation observed in mouse models of colitis^11,13^. This atypical nuclear receptor contains a well-ordered hormone-binding pocket, which binds signaling phospholipids including phosphoinositides^15,16,17,18^. However, structural and biochemical studies have revealed major differences between the human and mouse orthologues; hLRH-1 manifests a greater ligand-binding dependency^15,17,19^.

Here we investigate the physiological and pathophysiological function of hLRH-1 in the intestinal epithelium. Using humanized mouse intestinal organoids, a humanized in vivo IBD model, and human intestinal organoids, we uncover an essential role for LRH-1 in intestinal epithelial homeostasis and cell survival, which mitigates inflammatory injury. Our data rationalize the effort required to target this nuclear receptor for the treatment of IBD.

## RESULTS

### LRH-1 maintains epithelial integrity and viability

In order to investigate the role of LRH-1 in gut epithelia, LRH-1 expression and the effects of its deletion were determined in mouse intestinal organoids. Similar to prior in vivo studies^6^, *mLrh-1* was found in the crypt domain of intestinal organoids, but was also detected at lower levels in the villus domain (Figure 1a). Using *Lrh1^fl/fl^*; *VilCre-ERt2* (*Lrh1^IEC-KO^*) mice, intestinal organoids were generated following conditional and acute deletion of mLRH-1 (Figure 1b). Consistent with the proposed role for LRH-1 in Wnt/b-catenin-regulated cell growth^6^, deletion of mLRH-1 increased cell death and lowered organoid viability in a modified MTT reduction assay^20^, compared to control organoids from *Lrh1^fl/fl^* mice (Figure 1c).

**Fig. 1.**
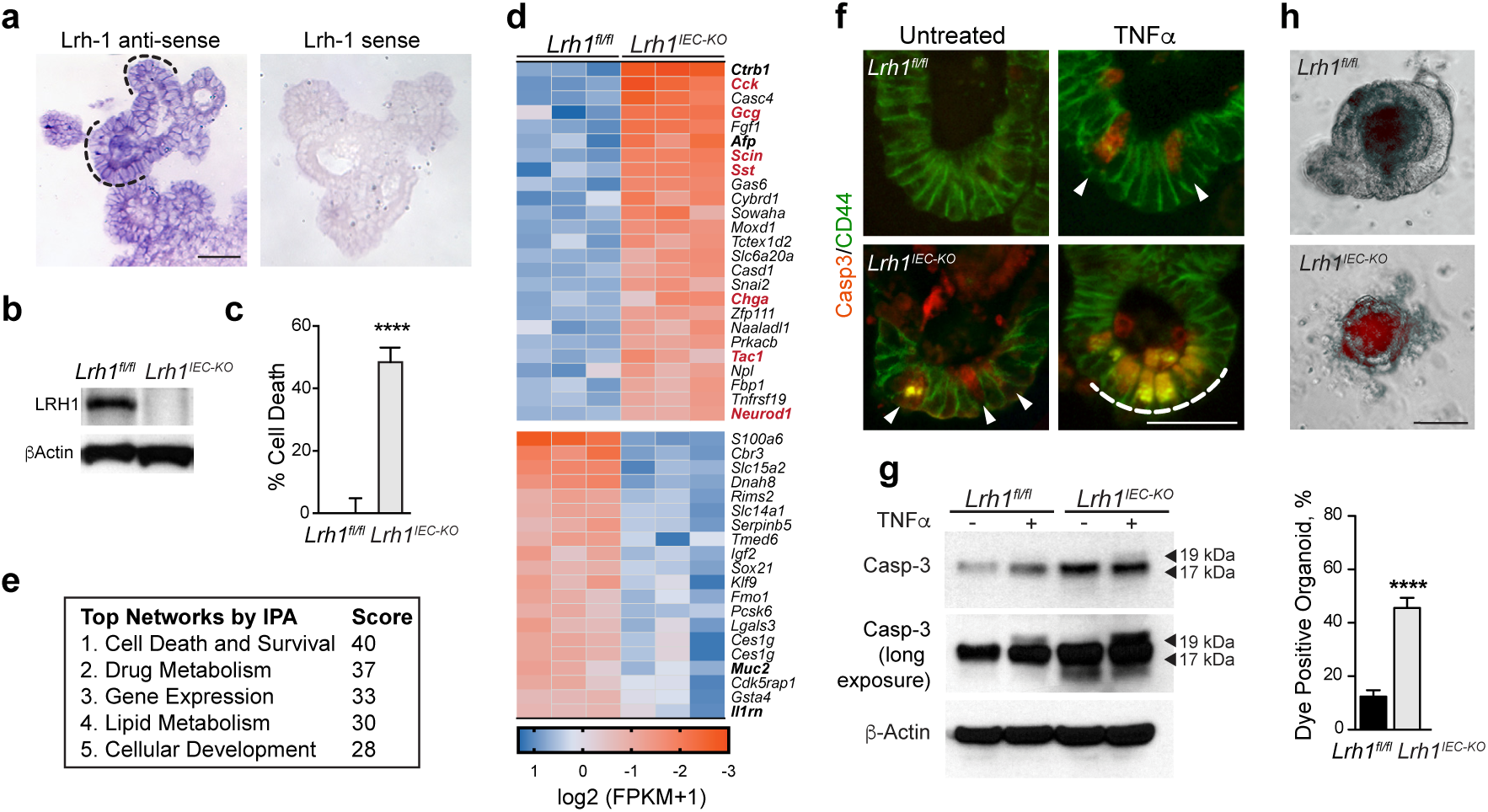
Loss of LRH-1 increases cell death and affects survival gene programs and epithelial permeability in murine organoids.++++**(a)** Expression of *mLRH-1* transcripts in intestinal organoids is widespread and higher in crypt regions (dotted black line). Scale bar = 100 mm. (**b)** Loss of mLRH-1 after Cre-recombination induced by 4-hydroxytamoxifen (4OHT) to *Lrh-1^fl/fl^;Vil-CreERT2* organoids for 48 hr (*Lrh1^IEC-KO^*) compared to similarly treated wild type (*Lrh1^fl/fl^*) organoids as detected by anti-LRH-1 antibody. **(c)** Cell death in *Lrh1^IEC-KO^* and *Lrh1^fl/fl^* organoids following acute loss of *Lrh-1*. Values normalized to 5 independent wells of untreated *Lrh1^fl/fl^* organoids taken to be 0%. **(d)** Most significant gene changes up (blue) or down (ochre) after loss of mLRH-1 by RNA expression (q = <0.05 and p = <0.005). Gene names in bold refer to markers of differentiation while red refer to EEC markers. **(e)** Top five altered gene networks as identified by IPA Ingenuity analysis. **(f)** *Lrh1^fl/fl^* and *Lrh1^IEC-KO^* organoids exposed to TNFα (10 ng/ml) for 40 h stained for active Casp3 (red) and CD44 (green). Cells expressing active Casp3 undergoing cell death are indicated (white arrowheads and dotted line). Scale bar = 50 µm. **(g)** Expression of active Casp3 in *Lrh1^fl/fl^* and *Lrh1^IEC-KO^* organoids as described in Panel F, detected by Western blotting (17kDa and19kDa, black arrowheads). Shorter (top panel) and longer exposures (middle panel). **(h)** Uptake of fluorescent dextran after 30 min in *Lrh1^fl/fl^* and *Lrh1^IEC-KO^* organoids. Scale bar = 100 µm. Bar graph of percentage dye-positive enteroids (≥ 30 total organoids counted per condition). Data represent average of minimum 3 biological triplicates. For **a** and **g**, Error bars are SEM using Student t-test (unpaired, 2-tailed) with p values of **** p = < 0.0001.

Transcriptional profiling of *Lrh1^fl/fl^* and *Lrh1^IEC-KO^* intestinal organoids revealed significant gene changes in cell survival and apoptosis pathways (Figure 1d, e), suggesting a role for LRH-1 in intestinal epithelial homeostasis and viability. Consistent with this notion, a marked increase in activated Caspase 3 (Casp-3) was observed in the intestinal crypt domain of *Lrh1^IEC-KO^* organoids, which was further exacerbated by TNFα (Figure 1f, g). As expected, given the documented role of LRH-1 in intestinal epithelial proliferation^6,21^, *Lrh1^IEC-KO^* intestinal organoids exhibited decreased cell proliferation measured by EdU incorporation (Supplementary Figure 1).

Because epithelial damage is a major contributor to chronic inflammatory disease in IBD^22^, we investigated whether loss of LRH-1 compromises the epithelial barrier. Indeed, significant failure of the epithelial barrier was observed in *Lrh1^IEC-KO^* intestinal organoids using a vital dye exclusion as-say (Figure 1h). Together, these data support an essential role for LRH-1 in epithelial viability and resilience.

### LRH-1 affects crypt survival and cellular differentiation via Notch signaling

Notch expression in the intestinal crypt preserves LGR5^+^ stem cells while restricting secretory lineages and is critical for ISC survival^23,24^. We then asked if the observed crypt cell death in *Lrh1^IEC-KO^* organoids might arise from impairment in Notch signaling. Indeed, both *Notch1* transcripts and protein levels were diminished in *Lrh1^IEC-KO^* organoids (Figure 2a,b). Because Notch is also a key factor in epithelial differentiation^23,24^, cell numbers and markers for Paneth, goblet, and enteroendocrine cells (EECs) were assessed after loss of LRH-1. As expected, lowered Notch signaling in *Lrh1^IEC-KO^* organoids resulted in downregulation of the stem cell markers *Lgr5* and *Olfm4*, while leading to upregulation of *Lys* and *Muc-2*; two respective markers for secretory Paneth and goblet cells (Figure 2b). The number of goblet cells doubled in *Lrh1^IEC-KO^* intestines, and Paneth cells were visibly expanded in intestinal crypts (Figure 2c, d). Surprisingly, rather than observing an expansion of EECs, as previously described with Notch inhibition^23,24^, the number of enterochromaffin cells, a representative population of EEC cells, was significantly reduced in *Lrh1^IEC-KO^* intestine (Figure 2c, d), as were levels of EEC-specific transcripts (Figure 1d). Collectively, these data imply that LRH-1 is necessary for maintenance of Notch signaling and cell survival and for proper allotment of intestinal epithelial cell types.

**Figure 2.**
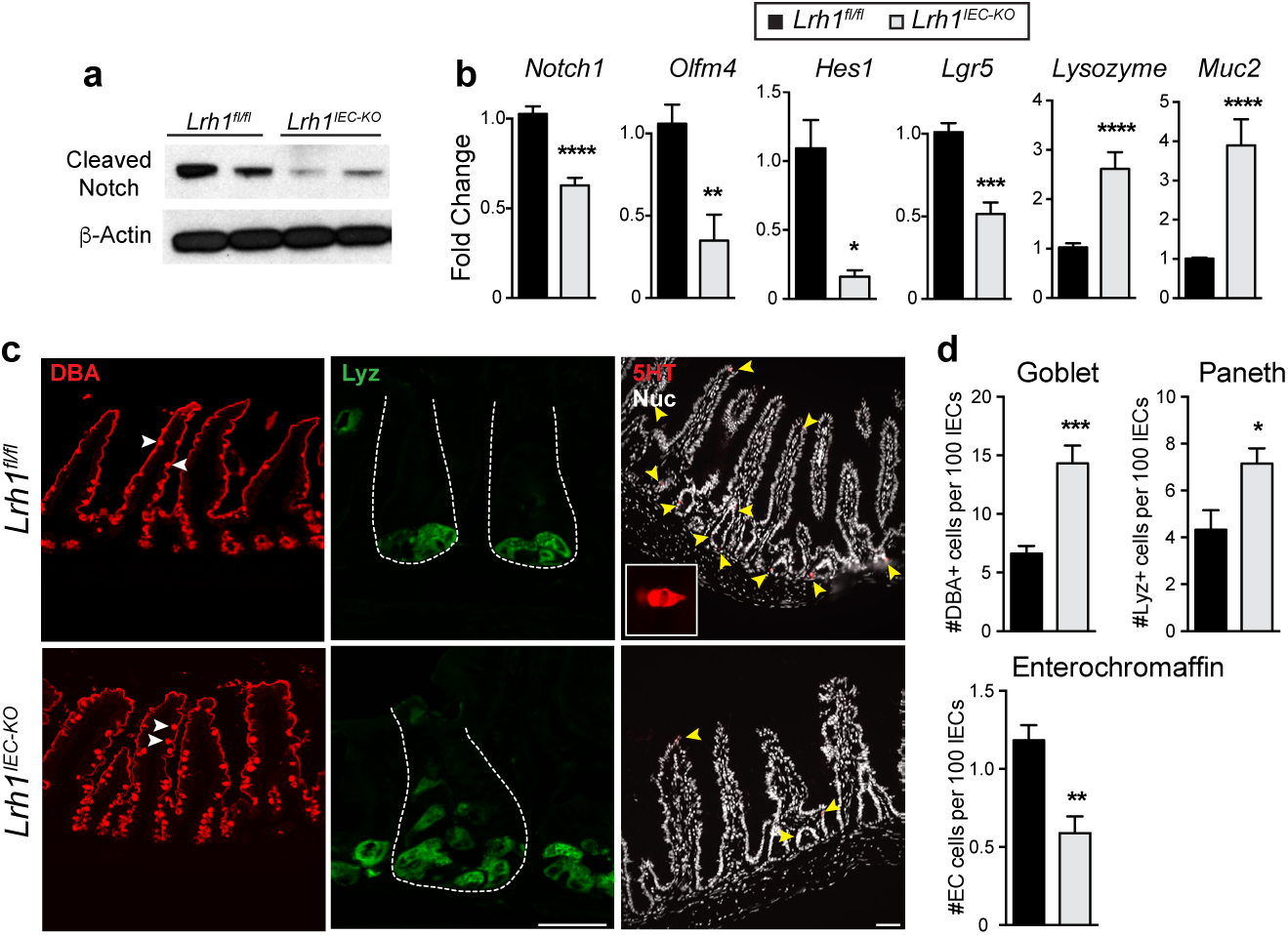
*Lrh1^IEC-KO^* mouse organoids have diminished Notch activation and undergo aberrant epithelial development.++++**(a)** Representative immunoblot for cleaved Notch1 in *Lrh1^fl/fl^* (left two lanes) and *Lrh1^IEC-KO^* (right two lanes), n = 4 per condition. **(b)** Fold change by RT-qPCR in *Lrh1^fl/fl^* (black) and *Lrh1^IEC-KO^* (grey) organoids for *Notch1* and Notch target genes *Olfm4* and *Hes1*. Stem cell marker *Lgr5*, Paneth cell marker *Lysozyme*, and goblet cell marker *Mucin2* are also shown. Minimum of 3 replicates per condition. **(c)** Histology of *Lrh1^fl/fl^* and *Lrh1^IEC-KO^* small intestine showing goblet (DBA, red, left), Paneth (lysozyme, green, middle) and enterochromaffin cells (5HT, red, right and inset). Scale bar = 50 µm. Intestinal crypts are outlined with white dashes in middle panel. **(d)** Quantitation of epithelial subtype distribution, n = 3 animals per condition. For **b** and **d**, error bars are SEM using Student t-test (unpaired, 2-tailed) with p values of * p = < 0.05, ** p = < 0.01, *** p = < 0.001, and **** p = < 0.0001.

### Human LRH-1 prevents crypt death and TNFα injury in mouse intestines

We next asked whether restoring or overexpressing LRH-1 might strengthen epithelial resilience to an inflammatory challenge. Human LRH-1 (hLRH-1), rather than mLRH-1, was chosen because it displays greater ligand-dependent activation and is the relevant isoform in human disease. As illustrated in Figure 3a, hLRH-1, unlike mLRH-1, lacks the salt-bridge at the mouth of the ligand binding pocket and requires a positive charge to stabilize this domain^15,25^. This role is fulfilled by the phosphate in the polar head group of its phospholipid ligand. Expression of hLRH-1 in mouse intestinal organoids was achieved by an AAV8-mediated infection protocol, which was optimized using AAV8-GFP. This method resulted in rapid and efficient gene expression (Figure 3b) that persisted for the life of the epithelial cell (Supplementary Figure 2) and permitted dosing that could either match or exceed endogenous levels of mLRH-1 (Figure 3c, left). Nuclear expression of hLRH-1 was detected throughout the crypt and villus zones (Figure 3b).

**Figure 3.**
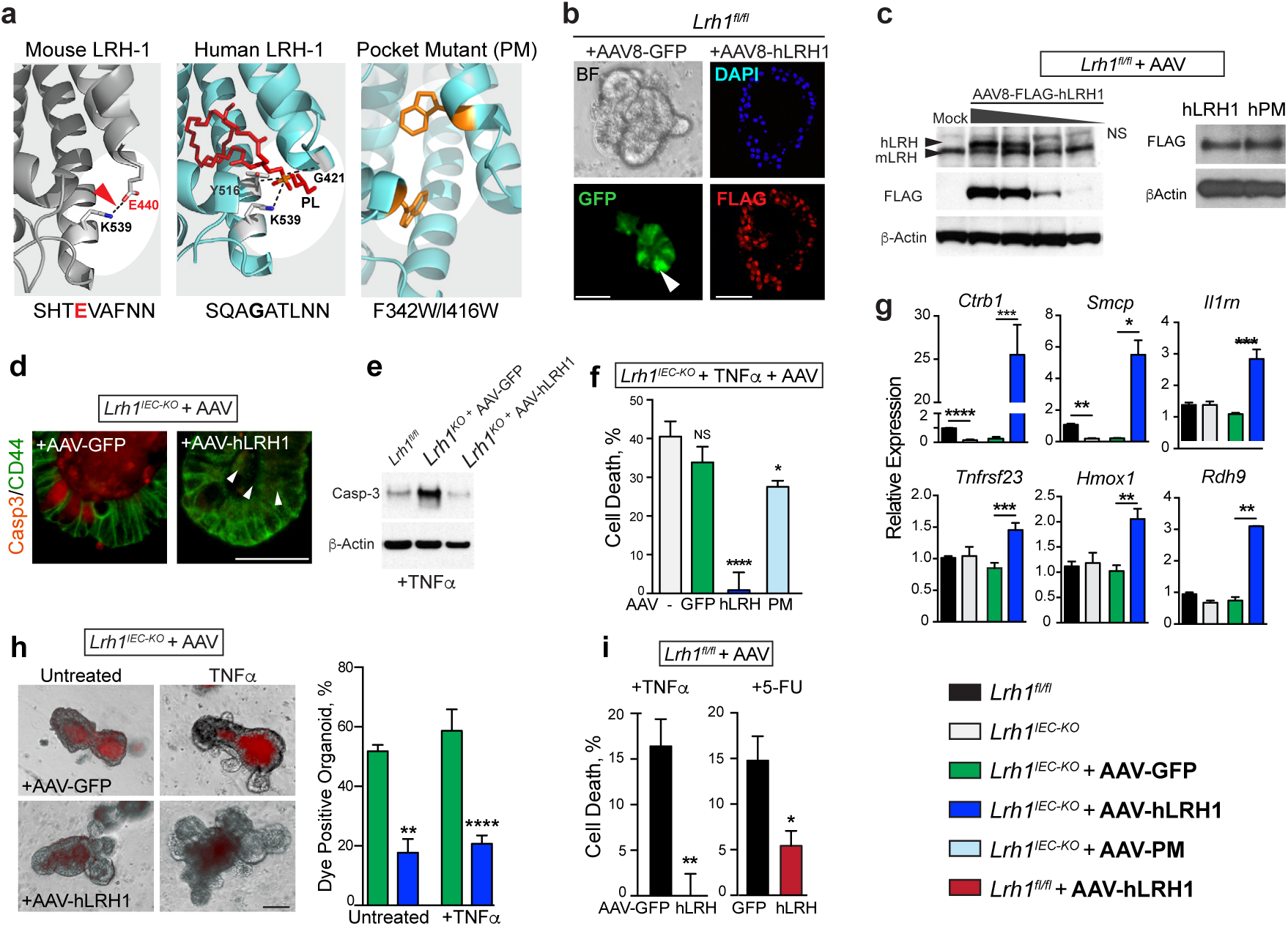
Rescue of *Lrh1^IEC-KO^* mouse organoids by hLRH-1 is ligand dependent and leads to an anti-inflammatory genetic program.++++**(a)** Ribbon diagrams of mouse (left) and human (middle) LRH-1 ligand binding pocket highlighting species-specific structural features of salt bridge (dotted black line and red arrowhead) coordination of phospholipid ligand (red stick; dotted black lines). Model of hLRH-1 pocket mutant (right) showing placement of pocket-obscuring residues (gold). **(b)** AAV8-directed GFP expression in organoids after 24 h (BF, brightfield; GFP, fluorescence; white arrowhead indicates representative GFP^+^ cell). Nuclear hLRH-1 expression 4 d post-infection detected with anti-Flag. Scale bar = 100 µm. **(c)** Titration of hLRH-1 protein by infectious titer of AAV8-hLRH1 (3.3 × 10^10^ – 4.1 × 10^9^ genome copies) after 4 d (left); mock infection is without virus. LRH-1 detected by anti-LRH-1 (upper panel) or anti-FLAG (middle panel). NS, non-specific band. Western blot for AAV-hLRH1 and AAV-PM detected by FLAG antibody from *Lrh1^fl/fl^* organoids infected with equal titer of AAV (right). **(d)** Casp3 signal in untreated *Lrh1^IEC-KO^* organoid crypts infected with either AAV-GFP or AAV-hLRH1 with Casp3^+^ cells indicated (white arrowheads). Scale bar = 50 µm. **(e)** Expression of active Casp3 protein in *Lrh1^IEC-KO^* organoids infected with AAV-GFP or AAV-hLRH1 for 72 h prior to TNFα (10 ng/ml, 40 h). **(f)** Percent cell death in *Lrh1^IEC-KO^* organoids infected with mock (grey), AAV-GFP (green), AAV-hLRH1 (blue), or AAV-hPM (hLRH-1 pocket mutant; light blue) for 72 h, and then treated with TNFα (10 ng/ml, 40 h). **(g)** Fold change by RT-qPCR in *Lrh1^fl/fl^* (black) and *Lrh1^IEC-KO^* (grey) organoids, or in *Lrh1^IEC-KO^* organoids subsequently infected with AAV-GFP (green) or AAV-hLRH1 (blue) for 72 h. **(h)** Uptake of fluorescent dextran in *Lrh1^IEC-KO^* organoids infected with AAV-GFP or AAV-hLRH1 for 72 h followed by TNFα (10 ng/ml, 40 h) as per Figure 1h. Scale bar = 100 µm. **(i)** Viability of TNFa-exposed *Lrh1^fl/fl^* organoids (20 ng/ml, 40h) and 5-FU (5 mg/ml, 24h), respectively, overexpressing hLRH-1 by approximately two times endogenous levels. Control organoids were infected with AAV-GFP (black bar). For Panels **f**-**i** error bars are SEM with statistical analyses as per Figure 1.

Human LRH-1 expression was able to fully rescue crypt cell death and maintain viability in *Lrh1^IEC-KO^* organoids even upon challenge by TNFα (Figure 3d-f). To ascertain if ligand binding is necessary for hLRH-1-mediated rescue, we next attempted to salvage organoid viability with the well-characterized ligand binding-defective variant of hLRH-1 (hPM; Figure 3a)^26,27^. Bulky hydrophobic residues were modeled in the binding pocket to impede ligand uptake without effecting protein integrity, as previously demonstrated in cultured cell lines (RE). Indeed, the hPM variant is stably expressed in intestinal organoids (Figure 3c, right), and despite the fact that the hPM retains modest transcriptional activity^26,27^, it failed to rescue TNFα-induced cell death in *Lrh1^IEC-KO^* intestinal organoids (Figure 3f, light blue bar).

Expressing hLRH-1 resulted in upregulation of known downstream targets including *Shp*, *Cyp11a1, and* Cyp11b1 as well as robust expression of new potential LRH-1 targets (*Ctrb1* and *Smcp*), and mediators of cell survival and anti-inflammatory responses, including the decoy receptors *Il1rn* and *Tnfrsf23* and the anti-apoptotic factor *Hmox1* (Figure 3g and Supplementary Figures 3 and 4). Adding hLRH-1 to *Lrh1^IEC-KO^* organoids also restored the integrity of the epithelial barrier, as demonstrated by a reduction of vital dye-positive organoids (Figure 3h). Moreover, increasing the dosage of hLRH-1 in the presence of wild type mLRH-1 ameliorated TNFα-induced cell death, suggesting that elevated LRH-1 activity protects against inflammatory damage (Figure 3i, left). This effect extends to other epithelial insults, as overexpression hLRH-1 also protected against damage by fluorouracil (5-FU), a common chemotherapeutic with intestinal toxicity (Figure 3i, right). Taken together, these data demonstrate that hLRH-1 fully substitutes for mLRH-1 to restore cell survival and activate anti-inflammatory programs.

To confirm the survival role of LRH-1 in vivo, we used a humanized intestinal mouse model in which mLRH-1 is deleted and hLRH-1 expressed in an inducible Cre-dependent manner (referred to as *hLRH1^IEC-Flex^*). Despite the lower protein levels of hLRH-1 as compared to endogenous mLRH-1 in *hLRH1^IEC-Flex^* organoids (Figure 4a), expressing hLRH-1 reduced cell death by nearly 50% and restored *Ctrb1* levels (Figure 4b-c). These ex-vivo results were confirmed in vivo by the near absence of cleaved Casp3 in intestinal crypts of the ileum (and to a lesser extent in villi) in *hLRH1^IEC-Flex^* mice compared to *Lrh1^IEC-KO^* (Figure 4d). Taken together, these data reveal that human LRH-1 can promote cell survival in murine intestinal epithelia lacking mLRH-1 and following challenge by TNFα.

**Figure 4.**
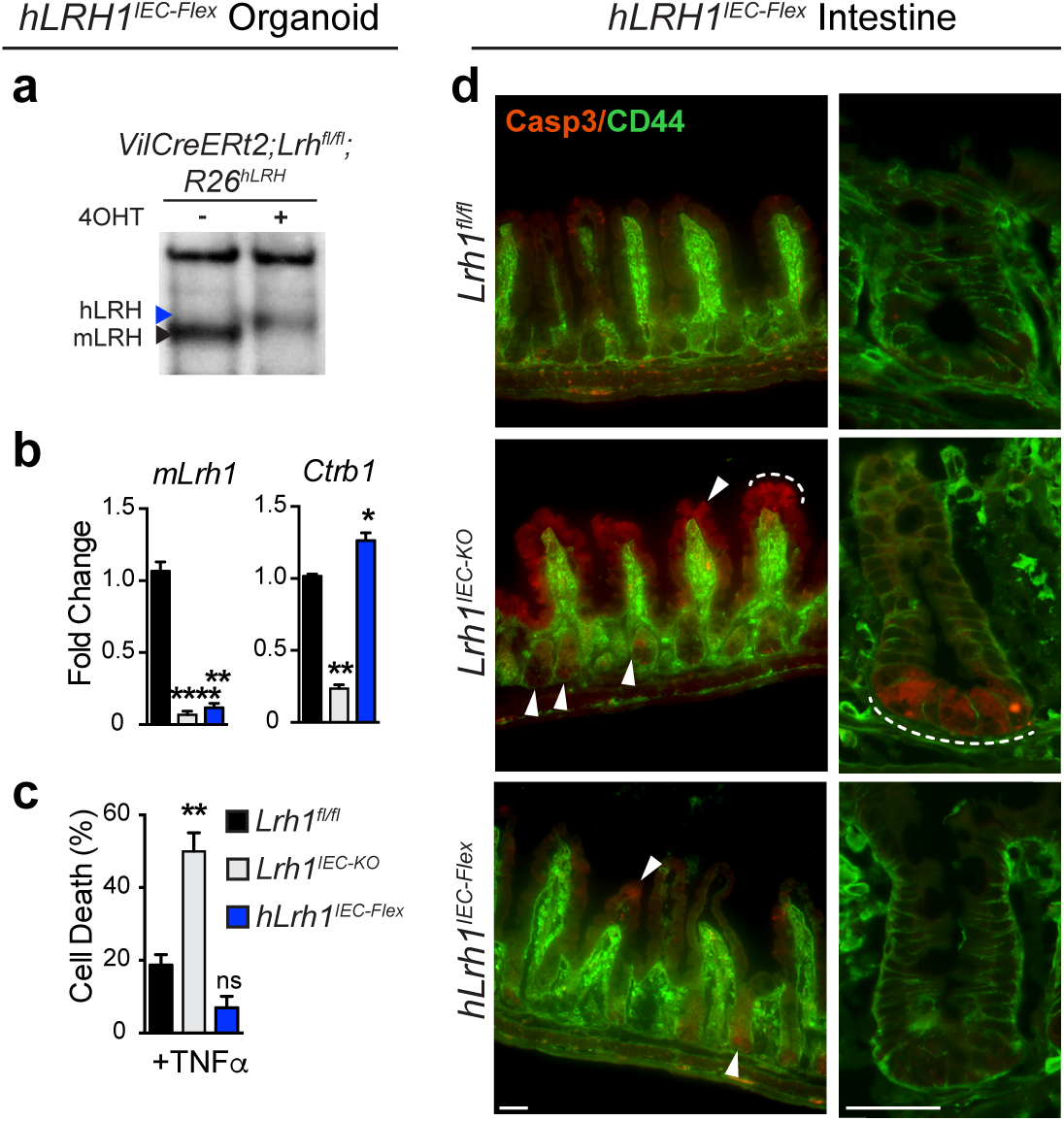
In vivo rescue of *Lrh1^IEC-KO^* mice by hLRH-1 reverses cell death.++++**(a)** LRH-1 protein levels in *hLrh1 ^IEC-Flex^* enteroids detected by anti-LRH-1 antibody with arrowheads indicating migration of human (blue) or mouse (black) LRH-1 before or after addition of 4OHT, which eliminates mLRH-1 and expresses hLRH-1; protein extracts were isolated 72 h later. **(b)** Relative levels of *mLRH-1* and a downstream LRH-1 target gene, *Ctrb1* in wild type (*Lrh1^fl/fl^*), *Lrh1^IEC-KO^*, and *hLrh1^IEC-Flex^* enteroids, with values normalized to wild type set at 1.0. Generation of *hLrh1^IEC-Flex^* is described in Materials and Methods. For all panels, data were generated from three independent wells of enteroids (~50 organoids per well) done in triplicate. **(c)** Percentage of cell death in *hLrh1^IEC-Flex^* enteroids with TNFα (10 ng/ml, 40 h) after eliminating mLRH-1 (grey) and expressing hLRH-1 (blue) by addition of 4OHT for 48 h. Data are also shown for treated *Lrh1^fl/fl^* enteroids (black). All values are normalized to 5 independent wells of untreated *Lrh1^fl/fl^* enteroids, which is taken to be 0%. **(d)** Immunofluorescence of wild type (*Lrh1^fl/fl^*), *Lrh1^IEC-KO^*, and *hLrh1^IEC-Flex^* ileum from adult male mice treated with two consecutive injections of tamoxifen. Staining for activated Casp3 (red) and CD44 (green), which marks intestinal epithelial crypt cells is shown at lower (first column) and higher (second column) magnification. The appearance of apoptotic cells is indicated in the crypt region as well as the villus (white arrowheads and dashed white line) in *Lrh1^IEC-KO^* ileum; some signal is also observed after expressing hLRH-1 in *hLrh1^IEC-Flex^*. Scale bars = 50 m. N = 2 per genotype. For Panels **b** and **c** error bars are SEM with statistical analyses determined by Student unpaired t-test, 2-tailed with p values of * p = < 0.05, ** p = < 0.01, and **** p = < 0.0001.

### hLRH-1 ameliorates immune-mediated colitis in vivo

To determine whether increased levels of LRH-1 can improve the course of disease in an immune-mediated model of colitis, *VilCre;Rag2^−/−^;Rosa26-Flox-Stop-Flox hLRH-1* mice (*Rag2^−/−^hLrh1^IEC-TG^*) were generated to conditionally overexpress hLRH-1 in the intestinal epithelium in the presence of endogenous mLRH-1. As expected from our earlier data, mLRH-1 elimination greatly exacerbated T cell transfer (TcT)-induced colitis in *Rag2^−/−^hLrh1^IKO^* male mice (Figure 5a). Significantly, this was markedly different after overexpressing hLRH-1. Disease severity was largely mitigated in *Rag2^−/−^hLrh1^IEC-TG^* mice, as evidenced by the relative preservation of body weight, prolonged disease-free survival, improved colitis histology scores and reduced disease activity index. In fact, nearly all disease parameters were better in *Rag2^−/−^hLrh1^IEC-TG^* animals relative to animals expressing endogenous mLRH-1 (Figure 5a). Mirroring this improvement in colitis, *Rag2^−/−^hLrh1^IEC-TG^* animals showed a decreased inflammatory cytokine profile, with lower intestinal expression of TNFα, IL-1b, and IL-6, and a corresponding increase in the anti-inflammatory cytokine IL-10 (Figure 5b). Collectively, these in vivo data establish the critical role of LRH-1 as an anti-inflammatory agent in the intestinal epithelium.

**Figure 5.**
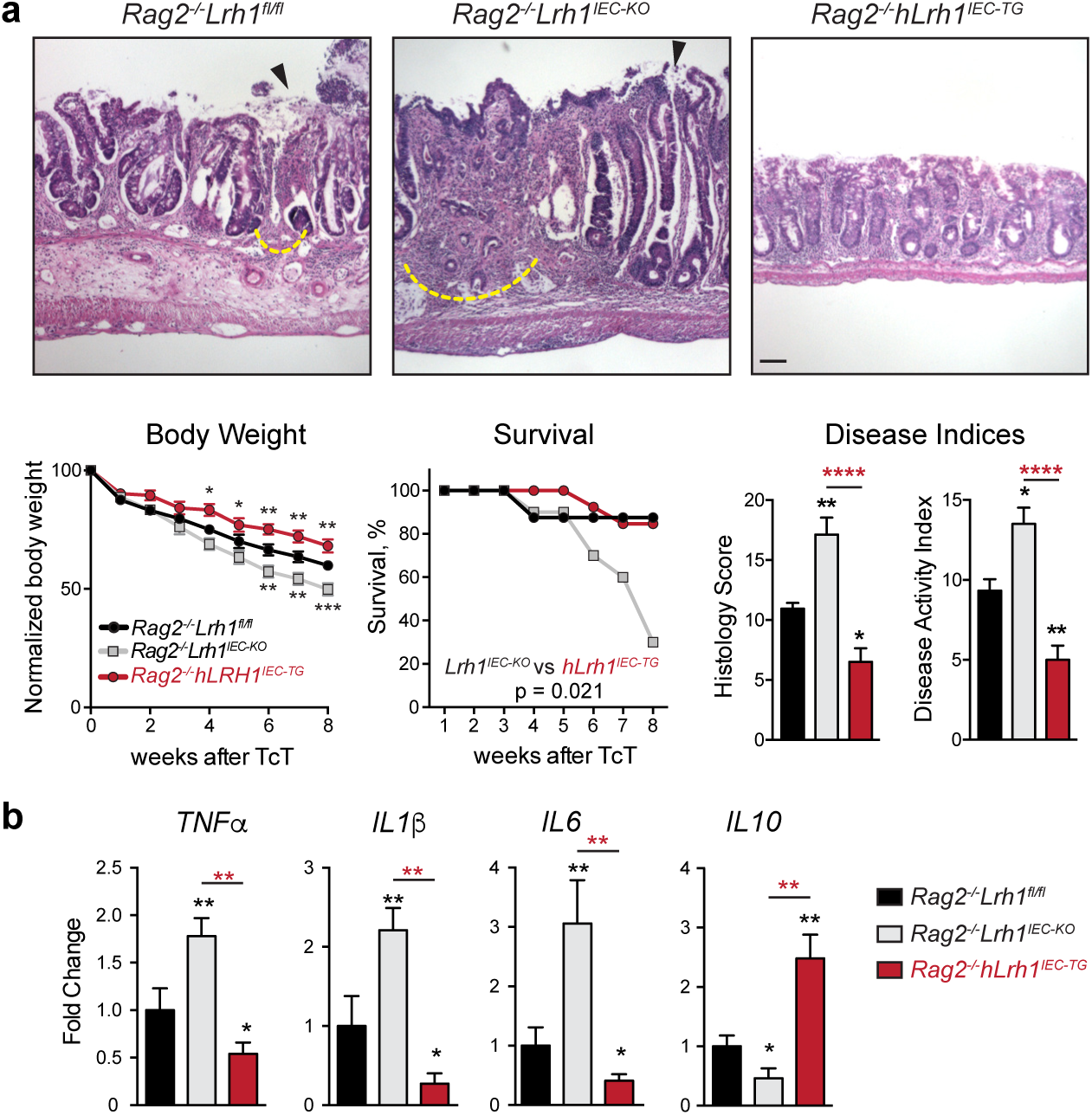
Overexpression of hLRH-1 decreases disease severity in a T cell transfer model of colitis. **(a)** Representative histology from T cell transfer colitis model for *Rag2^−/−^Lrh1^fl/fl^*, *Rag2^−/−^Lrh1^IEC-KO^*, and the LRH-1 overexpressing mouse line *Rag2^−/−^Lrh1^IEC-TG^*. Areas of mucosal erosion indicated with black arrowhead, crypt destruction indicated with dashed yellow line. Animal weight loss, survival during disease course, histology scores, and disease activity index are plotted below. Animal weight loss was normalized to non-diseased control at each time point. Scale bar = 100 µm. **(b)** Relative RNA levels of inflammatory and regulatory cytokines from colonic tissue of inflamed colon. Black and red asterisk symbols indicate comparison with *Rag2^−/−^Lrh1^fl/f^* and *Rag2^−/−^Lrh1^IEC-TG^* groups, respectively. Survival curves were determined by Kaplan-Meier survival analysis and Log-rank test; error bars are SEM. Statistical analyses for normalized body weights determined by two-way ANOVA and for histology and disease activity index determined by one-way ANOVA with p values of * p = < 0.05, ** p = < 0.01, and **** p = < 0.0001. For weight analysis, n = *Rag2^−/−^Lrh1^fl/fl^* no TcT (n = 4), *Rag2^−/−^Lrh1^IEC-KO^* (n = 6), *Rag2^−/−^Lrh1^IEC-TG^* (n = 6). For survival analysis, n = *Rag2^−/−^Lrh1^fl/fl^* (n = 8), *Rag2^−/−^Lrh1^IEC-KO^* (n = 10), *Rag2^−/−^Lrh1^IEC-TG^* (n = 13). For DAI, histology score and qPCR analysis, n=6 for each group.

### LRH-1 protects human intestinal organoids from TNFα injury

To determine whether the anti-inflammatory and pro-survival activity of LRH-1 translates to the human intestinal epithelium, human small intestinal organoids were derived from endoscopic biopsy samples of ileum from both Crohn disease patients and healthy individuals (Figure 6a). Unlike murine small intestinal organoid cultures, human derived organoids are maintained in a partially differentiated state, consisting of stem cells and partially differentiated Paneth cells, and undergo differentiation following WNT withdrawal^4^. Human *Lrh1* is expressed at similar levels in both differentiated and undifferentiated human organoids (Supplementary Figure 5), and remains broadly distributed throughout the epithelium following differentiation, as confirmed by staining for secretory goblet (MUC2) and Paneth (LYZ) cells (Figure 6b). This pattern closely matches the broad distribution of murine *Lrh1* (Figure 1a).

**Figure 6.**
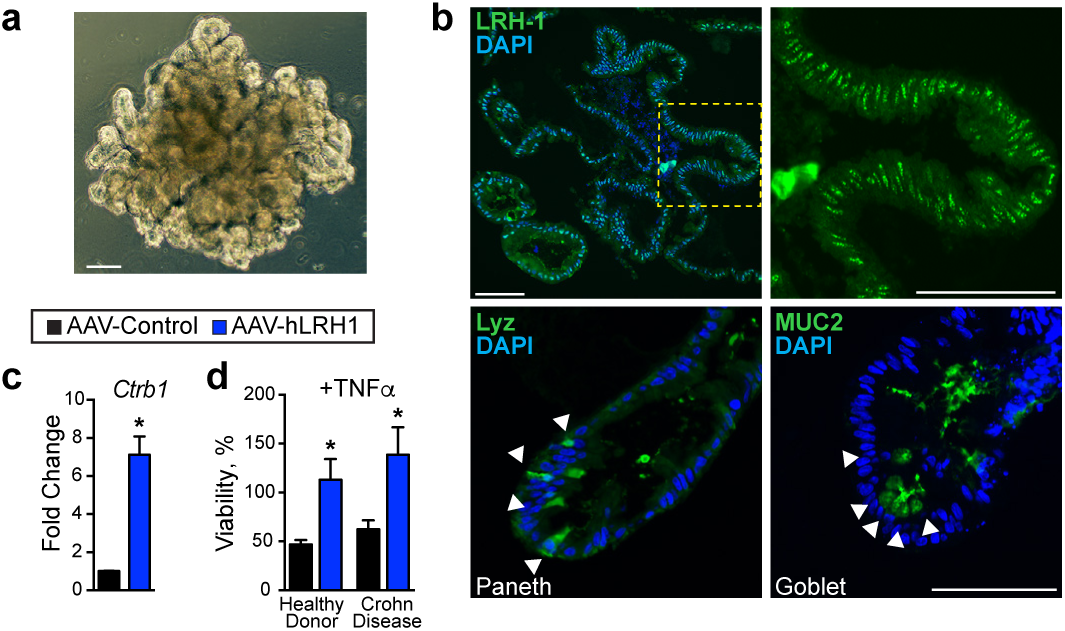
Increasing hLRH-1 levels in human intestinal organoids protects from TNFa-mediated cell death. **(a)** Bright-field view of human small intestinal organoid. Scale bar = 100 µm. **(b)** Immunofluorescence for LRH-1 (green, top panels) in human intestinal organoid sections shows expression throughout the organoid with strongest expression occurring in the crypt domain (yellow box and zoomed image, right). Differentiation markers for Paneth (Lyz, left) and goblet (Muc2, right) cells are shown below. Scale bar = 100 µm. **(c)** Expression of LRH-1 target gene *Ctrb1* in human intestinal organoids is upregulated 72 h after infection with AAV-hLRH1 but not a control AAV. **(d)** Over expression of hLRH-1 by AAV confers resistance to TNFa-mediated cell death. Human organoids from healthy donor and Crohn disease patient were infected with AAV-hLRH (blue) or AAV-Control (black) (3.3 × 10^10^ genome copies) for 72 h under differentiation conditions and then exposed to TNFa (20 ng/ml, 40 h). Data represent an N of at least three replicates with ~50 organoids per well. For Panels **c** and **d** error bars are SEM with statistical analyses determined by Student unpaired t-test, 2-tailed with p values of * p = < 0.05.

Increasing hLRH-1 dosage by AAV-mediated infection caused an upregulation of the LRH-1 target *Ctrb1* (Figure 6c). Importantly, in human organoids from both healthy individuals and Crohn disease patients, overexpression of hLRH-1 abrogated TNFα-induced cell death (Figure 6d). Taking all the data in this study together, we conclude that LRH-1 plays an essential role in intestinal homeostasis and in ameliorating inflammation-induced injury in human intestinal epithelia.

## DISCUSSION

In this study, using multiple independent mouse and human ex vivo and in vivo intestinal models, we establish that the nuclear receptor LRH-1 (*Nr5a2*) has a crucial role in maintaining the intestinal epithelium. Acutely knocking out mLRH-1 resulted in decreased Notch signaling and increased cell death in the intestinal crypt. Importantly, humanization of the mouse intestinal epithelium by expression of hLRH-1 corrected these deficits. Moreover, overexpression of hLRH-1 in both mouse and human intestinal organoids imparted epithelial resistance to both TNFα, a major inflammatory cytokine in inflammatory bowel disease, and 5-FU, a chemotherapeutic with intestinal toxicity. In the intact animal, expression of hLRH-1 ameliorated immune-mediated colitis. Using a viral-mediated approach newly applied to intestinal organoids, we showed that efficient rescue by hLRH-1 is ligand dependent. These findings are important because they provide a compelling argument that drug targeting of LRH-1 could enhance resistance to inflammation and restore intestinal epithelial health in intestinal diseases such as IBD.

Our study extends prior studies, reporting impaired cell renewal and enhanced chemical-induced colitis in heterozygous and conditional knockout mice, by demonstrating both a fundamental role for LRH-1 in the maintenance of epithelial viability and cell types, and the therapeutic potential of LRH-1 in intestinal disease. Further, we reveal that acute loss of mLRH-1 disrupts Notch expression, triggers increased apoptosis in the crypt and results in a breach in the epithelial barrier. Our data are consistent with the findings by Samuelson and colleagues that attenuating Notch signaling by genetic or pharmacological methods resulted in significant ISC apoptosis, crypt disruption, and expansion of secretory lineages. Interestingly, loss of LRH-1 not only recapitulates these findings, but also appears to compromise cells outside of the crypt base. We hypothesize that intestinal crypt apoptosis hinders renewal of the intestinal lining and exacerbates the immune inflammatory response3. In support of this idea, we demonstrate that loss of intestinal LRH-1 expression is associated with diminished animal survival and increased intestinal inflammation in the T cell transfer model of colitis. Importantly, increased hLRH-1 has a clear beneficial impact on disease activity and colitis scores, consistent with our organoid models.

The ability to acutely knock out mLRH-1 and replenish with hLRH-1 in intestinal organoids has provided new insights into the identity and function of species-specific LRH-1 targets in the small intestine. Based on the rapid crypt cell death and spectrum of differentially expressed genes, cell survival is the most prominent pathway affected following acute loss of mLRH-1. Interestingly, despite the known species difference in ligand binding^15^, hLRH-1 not only functionally complements mLRH-1 but also upregulates anti-inflammatory genes. hLRH-1 enhances *Il1rn* and *Tnfrsf23*, which act as decoy receptors for circulating pro-inflammatory cytokines. *Rdh9* (retinol dehydrogenase) was also upregulated 3-fold by hLRH-1; interestingly, is also increased in the intestine of conventional as opposed to germ-free mice^28^. *Ctrb1*, encoding chymotrypsin, and *Smcp* appear to be two highly sensitive markers of LRH-1 activity that are robustly activated by hLRH-1 in both mice and human intestinal organoids. It is known that LRH-1 binds the proximal promoter of *Ctrb1*^29^, raising the possibility that fecal chymotrypsin may serve as a biomarker to assess and follow pharmacological manipulation of hLRH-1 activity in vivo. Finally, ex vivo and in vivo mouse models of enteritis, in which mLRH-1 is replaced with the human form that binds signaling phospholipids more efficiently^15,26,30^, will provide a valuable platform to test and probe the utility of any synthetic ligands. In this regard, we note that the published LRH-1 agonist RJW100^30^ failed to activate the established and new gene targets that were identified in this study (data not shown).

Our data suggest that LRH-1 may play a previously unappreciated role in epithelial cell differentiation in the intestine, in addition to intestinal stem cell maintenance. Indeed, a similar role for LRH-1 has been reported in the pancreas^29,31^ and recently in neural stem cells^32^. Interestingly, although we observe an increase in secretory Paneth and goblet cells, consistent with reduction of Notch signaling, loss of LRH-1 also leads to a significant drop in markers defining nearly all subclasses of enteroendocrine cells^33^. These data infer separate but positive roles for LRH-1 in Notch intestinal crypt signaling and EEC differentiation. Intriguingly, the latter effect may be regional, with the greatest loss of EEC cells following deletion of LRH-1 in the ileum and proximal colon (J.B. and H.I., unpublished data); both of which exhibit high LRH-1 expression^6^. An understanding of how and where LRH-1 promotes lineage commitment in the gastrointestinal tract remains to be determined.

An important unanswered question is whether increased LRH-1 expression drives unchecked proliferation and promotes dysplasia in the intestinal epithelium, as suggested previously. Indeed, an earlier study showed that LRH-1 haploinsufficiency reduced tumor burden in the APCMIN model, possibly through interactions with the Wnt pathway at the *Cyclin D1* and *Cyclin E1* promoters^8^, but this same study noted decreased *Lrh1* in intestinal tumors. While our studies with the non-replicating AAV vector preclude us from exploring the question of cell proliferation in infected intestinal organoids, we note that neither *Cyclin D1* nor *Cyclin E1* were changed after loss of mLRH-1 or expression of hLRH-1 (Supplementary Figure 4).

The findings presented here leverage a new use for AAV-directed gene expression to rapidly manipulate small intestinal organoids. Positive features of AAV infection, versus classical lentiviral transduction, are high infectivity, rapid onset of gene expression, and the ability to infect large organoid fragments. On the other hand, the non-replicating nature of AAV restricts expression to the typical ~7 days turnover for mouse intestinal organoids. Nonetheless, this system allows a rapid structure-function analysis by simultaneously knocking out and adding back variants, which we put to effective use in this study to show the ligand-dependency of hLRH-1 effects. As unique molecular signatures of intestinal epithelial subtypes continue to emerge^33, 34^, engineering cell-specific promoters into the AAV-system should allow a more granular functional assessment of individual intestinal epithelial cell types.

In summary, our study of human intestinal organoids, humanized murine intestinal organoids, and a humanized murine IBD model show that LRH-1 promotes normal intestinal epithelial homeostasis and can be leveraged protectively against intestinal inflammation.

## MATERIALS & METHODS

### Mice and Study Approval

Animal studies were conducted in accordance with IACUC guidelines in strict accordance with the recommendations in the Guide for the Care and Use of Laboratory Animals of the National Institutes of Health. All animal studies and procedures were approved by the Baylor College of Medicine and UCSF Institutional Animal Care and Use Committees. All human studies were reviewed and approved by the UCSF Institutional Review Board (IRB 15-17763) and utilized tissue from de-identified donors who had consented to the study.

Animals were housed and bred in SPF facility. Inducible IEC knockout line was created by crossing animals harboring CreERT2 under control of the *villin* promoter with *Lrh1^fl/fl^* animals and bred to homozygosity. *Lrh1^IEC-Flex^* line was generated with Lox-STOP-Lox-hLRH1 animals, gift of DeMayo (Baylor College of Medicine) crossed into our *Lrh1^IEC-KO^* line. For animal knockout and activation studies, tamoxifen was dissolved in sunflower oil and delivered by two intraperitoneal injections 48 hours apart (1mg per day). Mice were sacrificed and tissue collected five days following the last tamoxifen dose. For the T cell transfer model, *Lrh1^fl/f^;VilCre* animals were bred with *Rag2^−/−^* (Jackson Laboratory) to homozygosity to generate *Rag2^−/−^ Lrh1^IEC-KO^* animals. Likewise, *Rosa26-Flox-Stop-Flox hLRH-1;VilCre* animals were crossed with *Rag2^−/−^* animals to generate *Rag2^−/−^ hLRH1^IEC-TG^* animals. Chronic enterocolitis was induced by T cell transfer of 0.5 million naive T cells per mouse, as described^35^. Briefly, wild type splenic CD4^+^CD45RB^high^ cells were isolated by MACS separation and flow cytometry cell sorting (Supplementary Figure 6), and then transferred by intra-peritoneal injection to *Rag2^−/−^* mice.

### Clinical disease activity index (DAI) and colon colitis scores

To assess the clinical disease activity index (DAI) body weight loss, diarrhea, guaiac-postive hematochezia, and appearance were monitored daily during the experiment. The DAI was determined according to a published scoring system^36^ (Supplementary Table 1). For colon histological analysis, the colon was divided into three segments (proximal third, middle third, and distal third). Each segment was embedded in paraffin, sectioned at 5μm, and stained with hematoxylin and eosin. Histological analysis was performed in the Cellular and Molecular Morphology Core of the Digestive Disease Center at Baylor College of Medicine. The sections were blindly scored using a standard histologic colitis score^37^. Three independent parameters were measured: severity of inflammation (0-3: none, slight, moderate, severe), depth of injury (0-3: none, mucosal, mucosal and submucosal, transmural), and crypt damage (0-4: none, basal one-third damaged, basal two-thirds damaged, only surface epithelium intact, entire crypt and epithelium lost). The score of each parameter was multiplied by a factor reflecting the percentage of tissue involvement (x1, 0-25%; x2, 26-50%; x3, 51-75%; x4.76-100%) averaged per colon.

### Crypt Cultures

Intestinal crypt cultures were derived from *Lrh1^fl/fl^*, *VilCreERT2;Lrh1^fl/fl^*, and *VilCreERT2;Lrh1^fl/fl^;Rosa^hLRH1^* six week old male mice. intestinal organoids were generated as previously described^5^. Briefly, the small intestine was isolated and flushed with ice cold phosphate buffered saline (PBS) and opened longitudinally. Villi were mechanically removed and the intestine cut into 1-2 µm pieces. Intestinal fragments were then incubated in an EDTA containing solution at 4 °C for 30 minutes. The intestinal fragment suspension was fractionated and crypt-containing fractions passed through a 70-mM cell strainer for plating in Matrigel. Crypt-Matrigel suspension was allowed to polymerize at 37 °C for 10 minutes. intestinal organoids were grown in base culture media (Advanced DMEM/F12 media, HEPES, GlutaMax, penicillin, and streptomycin) supplemented with growth factors (EGF, Noggin, R-spondin; Peprotech), B27 (Life Technologies), N2 (Life Technologies), and N-acetyl cysteine (NAC; Sigma). To activate genetic recombination, (Z)-4-hydroxytamoxifen (4OHT; Sigma) was added at 300 nM for 48 h.

### Human Intestinal Organoid Culture

Intestinal organoids were generated as previously described^4^. Briefly, endoscopic biopsy samples obtained from the terminal ileum were processed under a dissecting microscope to liberate intestinal crypts using EDTA chelation and mechanical disruption. Crypts were screened through a 100 µm filter, centrifuged, and suspended in ice cold Matrigel. The suspension was plated on pre-warmed cell culture plates. Following polymerization of Matrigel, propagation media (50% conditioned L-WRN, supplemented with human EGF (Peprotech), A-83-01 (Tocris), SB202190 (Sigma), Gastrin (Sigma), Nicotinamide (Sigma), B27 (Life Technologies), N2 (Life Technologies), GlutaMax (Life Technologies), and HEPES (Sigma) in F12 Advanced DMEM (Life Technologies)) was added. For the first 48 h of culture, CHIR99021 (Stemgent) and thiazovin (Stemgent) were added to support stem cell growth. To induce differentiation, media was replaced with differentiation media (base culture media supplemented with 10% R-Spondin conditioned media, human EGF (Peprotech), human Noggin (Peprotech), A-83-01, Gastrin, NAC, B27, and N2).

### Expression Analysis

Immunofluorescence (IF) and RNA *in situ* hybridization were performed on 5 µm cryosections using standard procedures. DIG-labeled (Roche) riboprobes were generated from pCRII-TOPO plasmid (ThermoFisher Sci) with mLRH-1 cDNA corresponding to bases 595-1683. Antibodies against hLRH-1 (1:200, Sigma HPA005455). FLAG (1:300, Sigma F7425), cleaved Caspase-3 (1:1000 (WB) and 1:400 (IF), Cell Signaling 5A1E), CD-44 (1:500, Tonbo 70-0441), lysozyme (1:200, DAKO EC 3.2.1.17) and MUC-2 (1:300, Santa Cruz Biotechnology sc-15334) were used with Alexa Fluor-conjugated secondary antibodies 1:300 (Millipore, Invitrogen). For Goblet staining, Rhodamine labeled Dolichos Biflorus Agglutinin (Vector Labs) was used at 1:200 dilution.

### RNA Isolation and PCR

Intestinal organoids were washed in ice-cold PBS and suspended in Trizol solution (Ambion). RNA was isolated with Direct-zol spin columns (Zymo Research). DNAse treated total RNA was used to generate cDNA using Superscript II (Invitrogen). Sybr green-based qPCR (Quanta) was performed on an Applied Biosystems Model 7900HT with primers as per Supplementary Table 2. The DDCt method was used for calculation of gene expression using *Gapdh* as reference. For RNA-Seq studies, RNA was isolated on Day 4 following AAV infection. Ovation RNA-Seq System V2 (NuGEN) was used to generate the cDNA library for sequencing on an Illumina HiSeq 4000. Data were analyzed using the GALAXY program suite^38^. Pathway analysis and annotations were performed with Ingenuity IPA (Qiagen) and Genecodis^39^, respectively. Data to be deposited in GEO. For colitis experiments, total RNA was isolated from snap-frozen colon tissues using Trizol Reagent (Invitrogen) and prepared for the cDNA with qScriptTM cDNA Synthesis Kit (Quanta Biosciences). Colonic gene expression was determined by qPCR using SYBR Green master (Kapa Biosystems Inc.). mRNA levels were normalized by the 36B4 gene expression. Pre-validated primers for qPCR were purchased from Qiagen (https://www.qiagen.com/geneglobe/default.aspx).

### Viability and Proliferation Assays

Murine intestinal organoids were plated in 10 ml Matrigel drops onto a pre-warmed 96 well cell culture plate and following polymerization 100 mL pre-warmed organoid growth media was added. Following Cre activation with 300 nM 4OH-tamoxifen, cultures were incubated with mTNFα for 40 hours. Viability was assessed by a modified 3-(4,5-dimethylthiazol-2-yl)-2,5-diphenyltetrazolium bromide (MTT) reduction assay as described^20^. Briefly, intestinal organoids were incubated with 500 mg/ml MTT for two hours at 37 °C. Media was aspirated and Matrigel dissolved in 2% SDS for 2 h at 37 °C with shaking. MTT was then solubilized in DMSO and absorbance measured at OD_562_. To normalize for crypt seeding and background drop-out, data were normalized by resazurin where indicated in the text. Here, intestinal organoids were incubated with resazurin (10 mg/ml) for six hours prior to administration of mTNFα. Media was removed and fluorescence measured (excitation 530 nm, emission 590 nm) and used to normalize MTT values^20^. Experiments were repeated a minimum of three times with five replicates per experiment. For human intestinal organoids, plates were set as above and organoids grown initially in propagation media for 24 h to establish organoids and then switched to differentiation media for the remainder of the experiment. Human TNFα was added after 72 h. Viability was determined by MTT 40 h after TNFα administration.

For proliferation studies, intestinal organoid cultures were incubated with 5-ethynyl-2-deoxyuridine (EdU, 10 mM) for two hours and then fixed in 4% paraformaldehyde. Click-It chemistry was performed as per manufacturer’s recommendations on 5 mm cryo-sections (Life Technologies). For 5-FU experiment, organoids were incubated with 5-FU (5_g/ml in DMSO; Millipore) for 24 hours and viability determined as above.

### Immunoblotting

Protein was isolated from intestinal organoids grown in 24 well culture plates in RIPA buffer containing protease inhibitors (Roche). Samples were processed in a Biorupter prior to gel loading. Antibodies include LRH-1 (R&D Systems PP-H2325-00), b-Actin (Ambion AM4302), FLAG (Sigma F1804), cleaved Caspase-3 (Cell Signaling 5A1E), and cleaved Notch1 (Cell Signaling D3B8).

### Dextran Exclusion Assay

Intestinal organoids were exposed to mTNFα for 24 hours and then incubated in 1 mg/ml Texas Red labeled dextran (average weight 10kDa; Life Technologies) for thirty minutes. Following incubation, excess dye was removed by serial washes with PBS and the plate imaged immediately. Wells were scored for fraction of dye-retaining intestinal organoids. 30-50 intestinal organoids were seeded per well. Eight wells per experiment were scored for each condition. Opened intestinal organoids were excluded from analysis. Results were validated by a blinded, independent observer.

### AAV-directed gene expression

AAV viral particles expressing hLRH1 or GFP under direction of the thyroxine-binding globulin (TBG) promoter were obtained from the University of Pennsylvania Viral Core. intestinal organoids were isolated in cold PBS, pelleted at 1000*g*, and resuspended in ice cold Matrigel. The mixture was added to chilled Eppendorf tubes containing virus on wet ice and then aliquoted immediately onto pre-warmed cell culture plates. After Matrigel was set, organoid growth media was added.

### Imaging

Live cell and intestinal organoid immunofluorescence imaging was performed on an Olympus IX51 microscope equipped with a DP71 imager. Mouse intestinal imaging was obtained on a Nikon Eclipse Ti equipped with a DS-Qi2 imager or an Olympus BX40 microscope with Magnafire imager.

## ACKNOWLEDGEMENTS

We thank Dr. Francesco DeMayo for his generous gift of hLRH-1 knock in mice, and Dr. Melvin Heyman, Liz Garnett, Emily Stekol, and Emma Canepa for assistance with human intestinal biopsy procurement. We thank S. A. Kliewer and D. J. Mangelsdorf (UT Southwestern Medical Center) for the gift of *Lrh1^f/f^* mice. We additionally thank Drs. Elena Sablin, Diego Miranda, and William Krause for helpful discussions and reading of the manuscript and the members of the D. D. M. laboratory for comments and additional support. We thank the support from Cytometry and Cell Sorting Core at Baylor College of Medicine and that from the Texas Medical Center Digestive Diseases Center (DDC).

## Funding

This work was supported by F32 CA163092 (JRB), K12 HD072222 (JRB), K08 DK106577 (JRB), T32 DK007762 (JRB, RN), UCSF Program for Breakthrough Research (ODK and RJF), UCSF Resource Allocation Program (JRB), Rainin Foundation, R01 DK099722, ADA #7-14-MI-08 (HAI), CIRM RN3-06525 (ODK), R01 DK085372 (DDM), the Alkek Foundation and the Robert R. P. Doherty Jr-Welch Chair in Science (DDM). This project was supported in part by the Cytometry and Cell Sorting Core at Baylor College of Medicine with funding from the NIH (P30 AI036211, P30 CA125123, and S10 RR024574).

## AUTHOR CONTRIBUTIONS

JRB, HW, RJF, ODK, DDM, and HAI conceived and designed the research. JRB, HW, RN, MS, and HSE performed experiments. JRB, HW, DDM and HAI performed the data analysis and wrote the manuscript. All authors reviewed the final manuscript.

## Supplementary Information

**Supplementary Figure 1.** *Lrh-1* loss is associated with decreased epithelial cell proliferation

**Supplementary Figure 2.** Time-course of AAV-directed GFP expression in intestinal organoids

**Supplementary Figure 3.** RNA-expression profiling of AAV-hLRH1 rescued organoids reveals increased survival and anti-inflammatory transcripts.

**Supplementary Figure 4.** Expression of LRH-1 gene targets.

**Supplementary Figure 5.** LRH-1 expression is relatively unchanged from undifferentiated to differentiated human small intestinal organoids.

**Supplementary Figure 6.** Gating strategy for FACS of CD4+ T cells

**Supplementary Table S1.** Colitis Disease Activity Score modified for T cell transfer mouse enterocolitis

**Supplementary Table S2.** RT-qPCR Primers used in this study

